# Carbon starvation induces the expression of PprB-regulated genes in *Pesudomonas aeruginosa*

**DOI:** 10.1101/639112

**Authors:** Congcong Wang, Wenhui Chen, Aiguo Xia, Rongrong Zhang, Yajia Huang, Shuai Yang, Lei Ni, Fan Jin

## Abstract

*Pseudomonas aeruginosa* can cause severe infections in humans. This bacteria often adopt a biofilm lifestyle that is hard to treat. In several previous studies, the PprA-PprB two-component system (TCS), which controls the expression of type IVb pili, BapA adhesin, and CupE fimbriae, was shown to be involved in biofilm formation. However, signals or environmental conditions that can trigger the PprA-PprB TCS are still unknown, and the molecular mechanisms of PprB-mediated biofilm formation are poorly characterized. Here we report that carbon starvation stress (CCS) can induce the expression of *pprB* and genes in the PprB regulon. The stress response sigma factor RpoS, rather than the two-component sensor PprA, was determined to mediate the induction of *pprB* transcription. We also observed a strong negative regulation of PprB to the transcription of itself. Further experiments showed that PprB overexpression greatly enhanced cell-cell adhesion (CCA) and cell-surface adhesion (CSA) in *P. aeruginosa*. Specially, under the background of PprB overexpression, both of the BapA adhesin and CupE fimbriae displayed positive effect on CCA and CSA, while the type IVb pili showed an unexpected negative effect on CCA and no effect on CSA. In addition, expression of the PprB regulon genes displayed significant increases in 3-day colony biofilms, indicating a possible carbon limitation state in these biofilms. The CSS-RpoS-PprB-Bap/Flp/CupE/Tad pathway identified in this study provides a new perspective on the process of biofilm formation under carbon-limited environments.

**IMPORTANCE:** Typically, determining the external signals that can trigger a regulatory system is crucial to understand the regulatory logic and inward function of that system. The PprA-PprB two-component system was reported to be involved in biofilm formation in *Pseudomonas aeruginosa*, but the signals that can trigger this system are unknown. In this study, we found that carbon starvation stress (CSS) can induce the transcription of *pprB* and genes in PprB regulon, through an RpoS dependent pathway. Increase of PprB expression leads to enhanced cell-cell and cell-surface adhesions in *P. aeruginosa,* both of which are dependent mainly on the Bap adhesin secretion system and partially on the CupE fimbriae. Our findings suggest that PprB reinforces the structure of biofilms under carbon-limited conditions, and the Bap secretion system and CupE fimbriae are two potential targets for biofilm treatment.

---

*Pseudomonas aeruginosa* is a ubiquitous opportunistic pathogen responsible for many human infections, especially the cystic fibrosis found in some immunocompromised individuals (1–3). In most cases of chronic infections, bacteria live in biofilm communities, and increasingly, they are becoming resistant to human immunity and antibiotic treatments (4–11). Cells in biofilms are typically embedded within a self-produced matrix of extracellular polymeric substances (EPSs) containing polysaccharides, proteins, lipids, and nucleic acids (12–16). Due to its key role in protecting the interior of the community from being killed by antibiotics or immune cells, the dense extracellular matrix has attracted substantial attentions. Numerous studies have pointed out the importance of two extracellular polysaccharides, Pel and Psl, in maintaining functional biofilm structures in *P. aeruginosa* (17–21). Whereas one previous research had identified a hyper-biofilm phenotype that was independent of Pel and Psl (22). Cells in this biofilm have been characterized by PprB overexpression, decreased Type III secretion, and increased drug susceptibility (22). PprB is a two-component response regulator that controls the transcription of numerous genes in *P. aeruginosa* (22, 23). Moreover, the *pprB* mutant strain has recently been reported to form a compromised biofilm in microfluidics systems (24). These preliminary results demonstrate that PprB and its downstream regulated proteins can dominate the formation of a new type of biofilm.

The PprB regulon contains multiple open reading frames, including genes encoding for type I secretion system substrates (*bapA*-*bapD*), CupE CU fimbriae, and type IVb pili, all of which are positively and directly regulated by PprB at the transcriptional level (22). The predetermined *bapA*, *bapB*, *bapC*, and *bapD* (*PA1874-1877*) genes consist of an operon in which *bapA* encodes a large externalized repeat-rich protein considered to be an adhesin. In addition, BapA protein was found mainly in the classical supernant of bacterial culture and associated loosely with the cell surface (22). This raises the confusion about whether BapA can enhance cell adhesion to surfaces. Meanwhile, the CupE fimbriae is a cell-surface-associated organelle that plays an important role in both the microcolony and 3D mushroom formations during biofilm development (25). The type IVb pili is referred to as the tight adherence (Tad) pili and is important in bronchial epithelial cell adhesion and host-colonization (26, 27). In *P. aeruginosa*, the Flp pilin consists of the main structure of the type IVb pili fibre and the *tad* locus proteins (RcpC-TadG) are responsible for ordered secretion, folding and the assembly of tens of thousands of pilin subunits (28). Previous study had revealed that BapA adhesin, CupE fimbriae and type IVb pili together contribute to the aforementioned hyper-biofilm phenotype (22).

The phenotypes of PprB overexpression in *P. aeruginosa* have been well documented. However, in the wild type strain, the exact external signals or environmental conditions that trigger the PprB pathway remain unknown. Transcriptional studies of *flp*, *rcpC* and *cupE* promoters in shaking conditions have indicated that all of these genes are commonly inducted after cells entered the stationary phase (25, 26). In this study, we demonstrated that carbon starvation stress (CSS) can trigger the expression of multiple PprB-regulated genes in *P. aeruginosa*.

The induction of PprB-regulated genes is dependent on the RpoS-controlled overexpression of PprB, rather than on the signal transduction of the putative sensor kinase PprA. We further demonstrated the roles of type IVb pili, CupE fimbriae and BapA adhesin in the cell-cell adhesion (CCA) and cell-surface adhesion (CSA) of *P. aeruginosa*. We also observed significant transcriptional increases in PprB regulon genes in colony biofilms after 3 days of cultivation. The CSS-RpoS-PprB-BapA/Flp/CupE signaling pathway determined in this study provides a new perspective on the process of biofilm formation and may be helpful in directing biofilm treatment.

## Results

### *flp* transcription is induced under CSS

Using the super-folder green fluorescent protein (sfGFP), a reporter expression system (see Materials and Methods) was established to assess the transcriptional activity of *flp* promoter. The reporter strain was first cultured to exponential phase using sodium succinate as the sole carbon source. Then cells were washed and introduced to the same media without sodium succinate. *flp* expression responded quickly to carbon deprivation, with great heterogeneity between cells (Fig. 1A). The fluorescence showed an approximately 50-fold (P < 0.001) induction after 5 hours of carbon deprivation (Fig. 1B). To test whether carbon limitation is the only inducement for the induction of *flp* transcription, a bacterial culture that had already experienced 4 hours of carbon deprivation was supplemented with 30 mM sodium succinate. SfGFP fluorescence decreased quickly upon succinate addition, and the half-life period (1 hour) of this decay was approximately the doubling time of cells, indicating that gene transcription had halted immediately (Fig. 1B). Furthermore, *flp* transcription showed similar inductions when we replaced sodium succinate with other types of carbon sources and repeated the carbon deprivation experiment (Fig. 1C). All these results indicate that CSS can induce *flp* transcription in *P. aeruginosa*.

**Figure 1:**
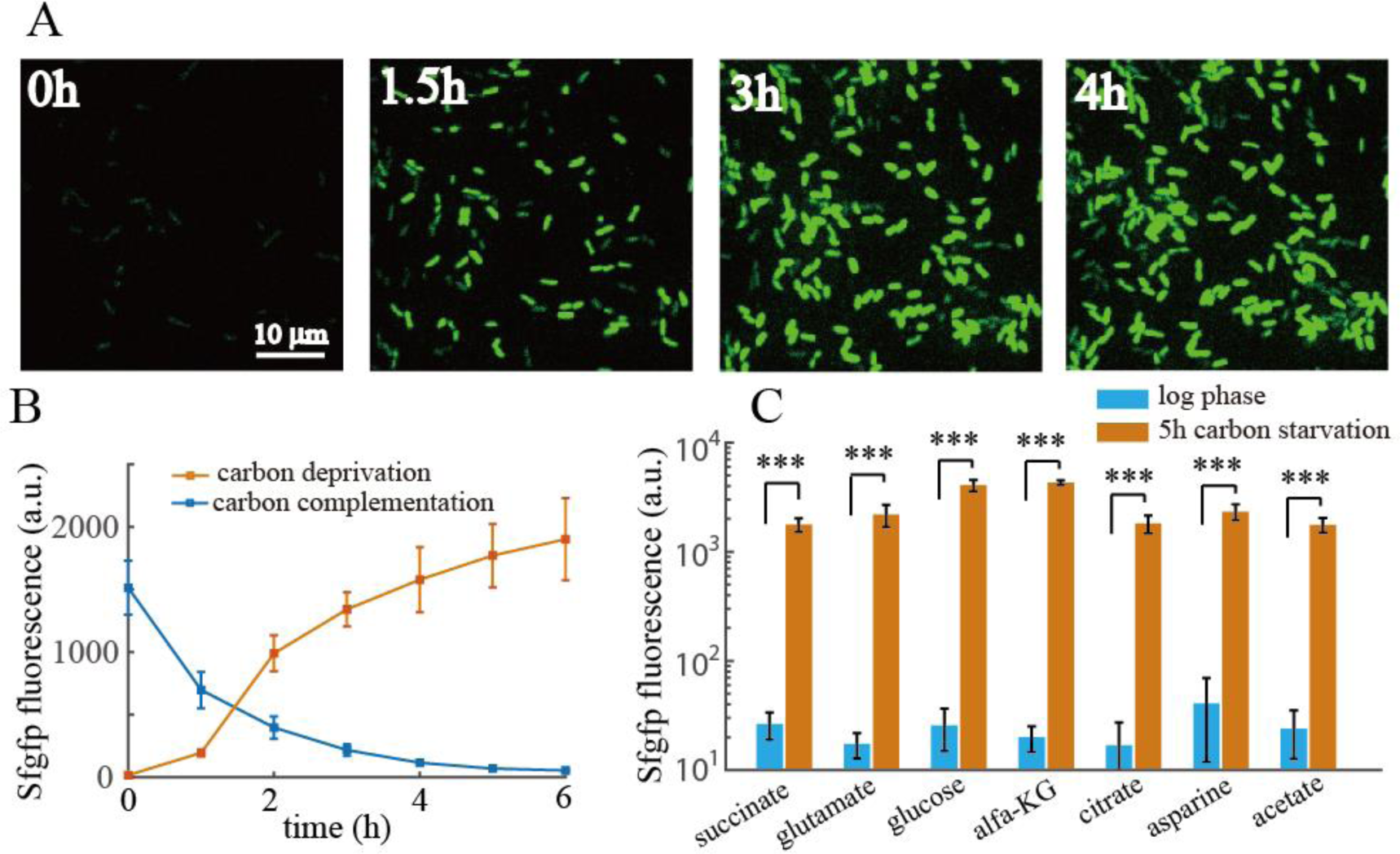
*flp* transcription is induced under CSS. (A), Sfgfp time-lapse imaging of *flp* transcriptional reporter cells after carbon deprivation. (B), Resulting expression values of *flp* transcriptional reporter over time after carbon deprivation (blue line) or *flp* expression over time after carbon complementation of 4-hour CSS pretreated cells (orange line). (C), Expression values of *flp* transcriptional reporter using different type of carbon sources at logarithmic phase or after 5-hour carbon deprivation. Statistical analysis used pairwise strain comparisons (t-test). ***P < 0.001.

### PprB is essential for the CSS response of *flp* transcription

We next investigated the potential regulators involved in the transcription of *flp* under CSS. Previously, *flp* transcription was reported to be mainly dependent on the PprA-PprB two-component regulatory system (26). Moreover, the carbon catabolite control system CbrAB-Crc-CrcZ in *P. aeruginosa* were found to be involved in the hierarchical management of carbon sources through regulation of gene expressions at both the transcriptional and translational level (29, 30). In addition, a LasR binding site had been predicted upstream of the *flp* coding sequence, suggesting that the quorum sensing system may also be involved in the control of *flp* transcription. We thus monitored the expression of *flp* reporter in *pprB*, *cbrAcbrB* and *lasRrhlR* mutant strains before and after carbon deprivation. *flp* expression in the *pprB* mutant completely lost the ability to respond to CSS, while in *cbrAB* and *lasRrhlR* mutants it displayed similar responses upon CSS to that of the wild type strain (Fig. 2A). Thus we concluded that PprB is essential for the CSS-induced expression of *flp*.

**Figure 2:**
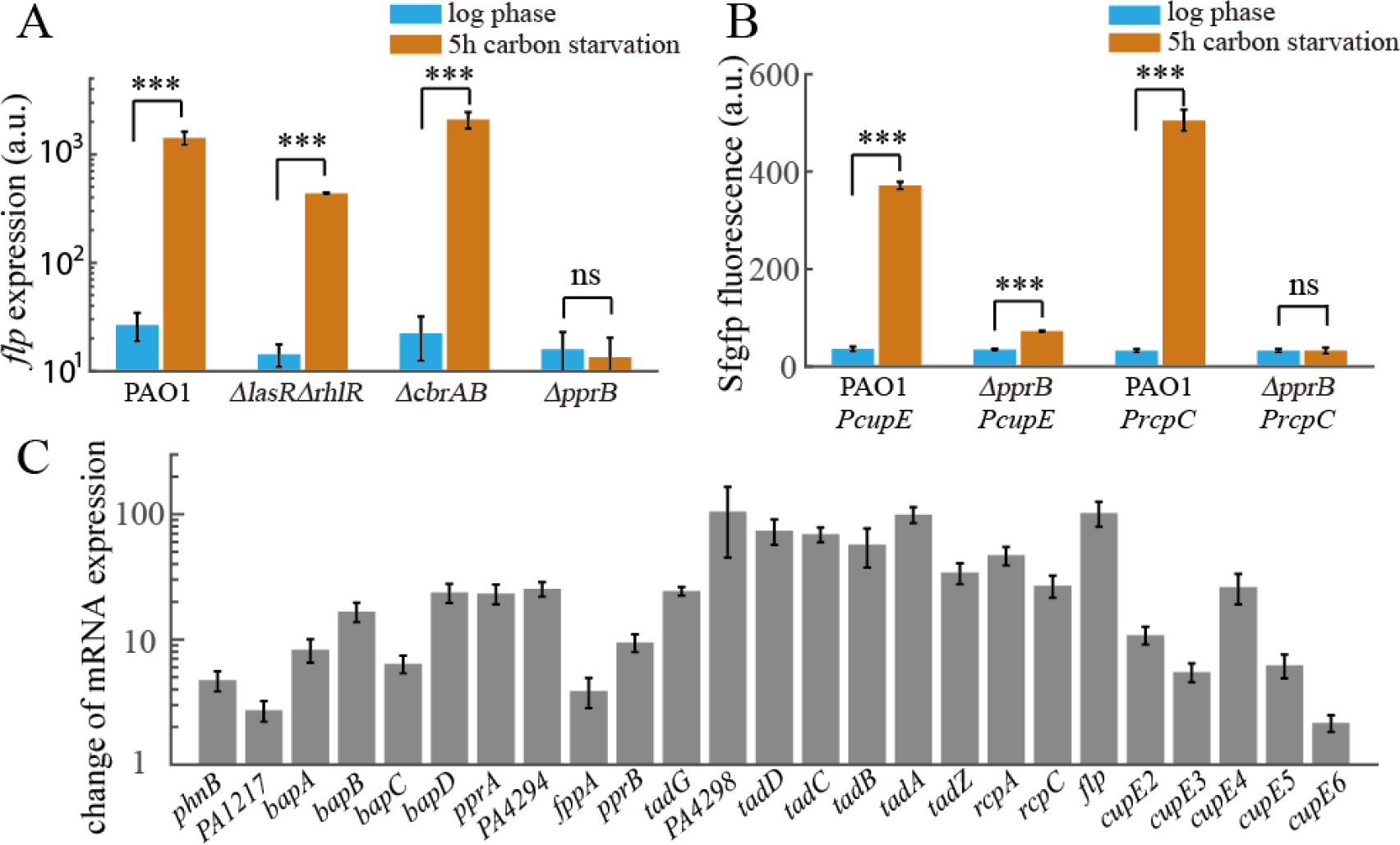
PprB-regulated genes are induced under CSS. (A), Expression values of *flp* transcriptional reporter in different mutants of *P. aeruginosa* at logarithmic phase or after 5-hour carbon deprivation. (B). Expression values of *cupE* or *rcpC* transcriptional reporters in wild type or *pprB* mutant strains at logarithmic phase or after 5-hour carbon deprivation. (C). RNA-seq fold change values of mRNA levels of PprB-regulated genes in response to CSS. All data are from three independent experiments and shown as the mean ± s.d. Statistical analysis used pairwise strain comparisons (t-test). ***P < 0.001; ns, non-significant.

### Transcription of *cupE* and *tad* locus are also induced under CSS and are PprB-dependent

Expression of two gene clusters, the *tad* locus encoding proteins required for type IVb pili assembly and the *cupE* locus encoding non-archetypal fimbrial subunits, are both controlled by PprB through direct transcriptional regulation (25, 26). We speculated that expressions of these two genetic loci are also upregulated under CSS. The fluorescent intensities of transcriptional reporters for *cupE* and *rcpC* were monitored in both wild type and *pprB* mutant strains. Consistent with our speculation, expression of *cupE* and *rcpC* showed 9- (P < 0.001) and 14- (P < 0.001) fold increase after carbon deprivation, the knockout of *pprB* eliminated the CSS response of *rcpC* and reduced the CSS response of *cupE* to 2 fold (P < 0.001) (Fig. 2B).

The PprB regulon had been previously determined (22), including genes involved in *Pseudomonas* quinolone signal (PQS) systems, type 1 secretion systems containing *bapA*, *bapB*, *bapC*, and *bapD* and the aforementioned type IVb pili and CupE fimbriae assembly systems. We further checked the responses of other PprB-regulated genes under CSS using RNAseq. Most of the genes within the PprB regulon were upregulated after 6 hours of carbon deprivation, with a fold change from 2 to nearly 100 (Fig. 2C). Therefore, the PprA-PprB two-component system was determined to be a key node during CSS response in *P. aeruginosa*.

### Increased expression of PprB under CSS contributes primarily to the transcriptional induction of PprB regulated genes

We then focused on the question of how the PprA-PprB system responds to CSS. In one previous study, PprA was thought to act as a sensor kinase, which is responsible for PprB phosphorylation after responding to external signals (23). Thus we naturally considered that the CSS signal should be transmitted through PprA. However, reporters in *pprA* mutant exhibited similar responses to CSS compared with those in the wild type strain (Fig. 3A, Fig. 3D). Therefore, although PprB dominates the transcriptional induction of related genes, CSS signal is not transmitted through PprA. Another possibility is that the CSS signal may trigger the expression of *pprB* (which was already observed in the RNAseq result), thereby up-regulating the expression of related genes. We further constructed a *pprB* transcriptional reporter in *P. aeruginosa* and investigated its response to CSS. Consistent with the RNAseq data, the fluorescence of the *pprB* reporter in wild type cells displayed approximately 10-fold increase (P < 0.001) upon 5 hours of carbon starvation (Fig. 3B).

**Figure 3:**
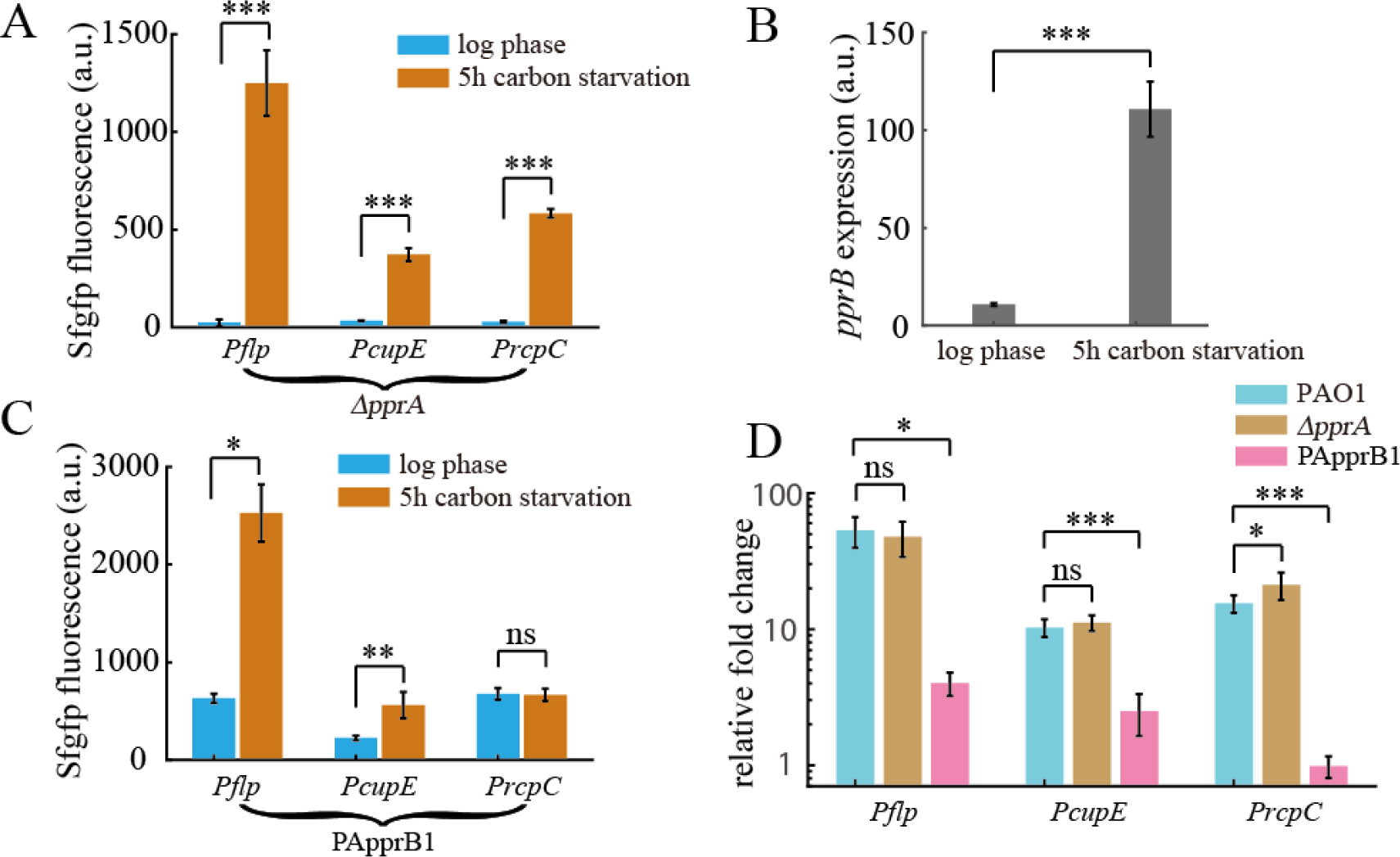
Increased expression of PprB under CSS contributes primarily to the transcriptional induction of PprB-regulated genes. (A), Expression values of *flp*, *cupE* or *rcpC* transcriptional reporters in *pprA* mutant strain at logarithmic phase or after 5-hour carbon deprivation. (B), Expression values of *pprB* transcriptional reporters in wild type strain at logarithmic phase or after 5-hour carbon deprivation. (C), Expression values of *flp*, *cupE* or *rcpC* transcriptional reporters in PApprB1 (PprB was constitutively overexpressed) strain at logarithmic phase or after 5-hour carbon deprivation. (D), Fold change values of *flp*, *cupE* or *rcpC* expression upon 5-hour carbon deprivation in wild type, *pprA* mutant or PApprB1 strains. All data are from three independent experiments and shown as the mean ± s.d. Statistical analysis used pairwise strain comparisons (t-test). *P < 0.05; **P < 0.01; ***P < 0.001; ns, non-significant.

To further check whether the transcriptional inductions of PprB regulated genes under CSS could be explained by the increase of PprB expression, reporters of *flp, cupE*, and *rcpC* were measured in a PApprB1 strain in which *pprB* was constitutively overexpressed. This PApprB1 strain was constructed by introducing the *pprB* gene into the chromosomal attTn7 site of the *pprB* knockout strain. Notably, the exogenously introduced *pprB* gene was driven by the arabinose inducible promoter P_BAD_. Expressions of the reporters at logarithmic phase increased more than ten folds in PApprB1 compared to that in wild type (Fig. 2B, Fig. 3C), consistent with the fact that PprB positively controls the transcription of these genes. However, CSS failed to induce the same expression changes of *flp*, *cupE and rcpC* in PApprB1. Only 3- (P < 0.05), 1.5- (P <0.01), and 0-fold increase for *flp*, *cupE and rcpC* were observed in PApprB1 (Fig. 3C, Fig. 3D), in contrast to the corresponding 50- (P < 0.001), 9- (P < 0.001) and 14-fold (P < 0.001) increase in PAO1 (Fig. 3D). Therefore, we concluded that the transcriptional inductions of *cupE*, *rcpC* and *flp* under CSS are primarily driven by the increased expression of PprB.

### Increased expression of PprB under CSS is controlled by RpoS

We then continued to search for the regulators that control the expression of PprB. In *P. aeruginosa*, the stress response sigma factor RpoS has been reported to enhance carbon starvation tolerance of bacteria (31, 32), thus, it is reasonable to speculate that RpoS is involved in the regulation of genes with altered expressions during CSS. In the *rpoS* mutant, Sfgfp fluorescence for *pprB* transcriptional reporter was barely detectable under CSS, complementing the *rpoS* mutation in PArpoS restored Sfgfp fluorescence (Fig. 4B), indicating the pivotal role that RpoS plays for *pprB* transcription. Based on the previously identified RpoS-dependent promoter consensus (33), a putative RpoS binding site (CTATATG) was mapped in the *pprB* promoter sequence (Fig. 4A). The *PpprB-mut1* reporter, whose RpoS binding site was mutated (CTATATG to GGGTATG), also failed to respond to CSS in the wild type strain (Fig. 4B). We also monitored the expression of *pprB* under the treatment of nitrogen starvation or acetate stress in which the RpoS activity could also be induced (34, 35), over 10-fold (nitrogen starvation, P < 0.001) and 12-fold (acetate stress, P < 0.001) increase were observed (Fig. 4C). These results strongly suggest that RpoS directly regulates the transcription of *pprB* and mediates the CSS response on *pprB* expression.

**Figure 4:**
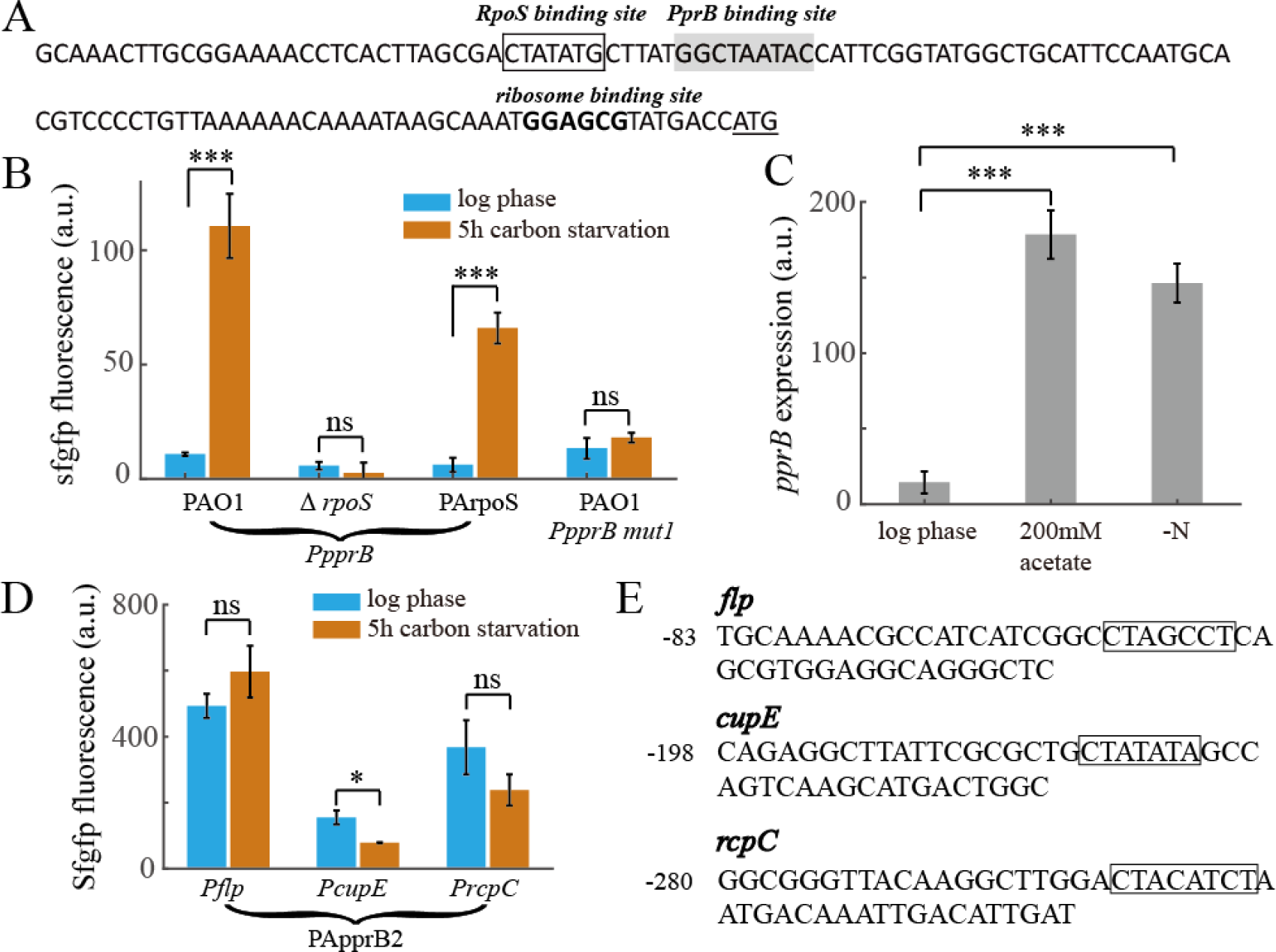
Increased expression of PprB under CSS is controlled by RpoS. (A), Promoter region of the *pprB* gene. Putative RpoS or PprB binding sites are framed or greyed. The ribosome binding site is shown in boldface and the translational start codon is underlined. (B), Expression values of *pprB*, or *PpprB mut1* (RpoS binding sequence CTATATG was mutated to GGGTATG) transcriptional reporters in the wild type or *rpoS* mutant or PArpoS (Δ*rpoS, rpoS* complement at genomic attTn7 site) strains at logarithmic phase or after 5-hour carbon deprivation. (C), Expression values of *pprB* transcriptional reporters in wild type strain at logarithmic phase or after 5-hour 200 mM acetate stress or 5-hour nitrogen starvation stress. (D), Expression values of *flp*, *cupE* or *rcpC* transcriptional reporters in PApprB2 (Δ*rpoS,* PprB overexpression) strain at logarithmic phase or after 5-hour carbon deprivation. (E), Promoter regions of the *cupE, rcpC and flp* genes. Putative RpoS binding sites are framed. Data in B, C, and D are from three independent experiments and shown as the mean ± s.d. Statistical analysis used pairwise strain comparisons (t-test). *P < 0.05; ***P < 0.001; ns, non-significant.

Expressions of *flp*, *cupE* and *rcpC* were further examined in strain PApprB2, in which *rpoS* was knocked out and *pprB* was overexpressed by the P_BAD_ promoter. All the three reporters in the PApprB2 strain displayed decreased expression with respect to that in PApprB1, especially under CSS, and the transcriptional induction of *flp* and *cupE* found in PApprB1 under CSS are completely abrogated (Fig. 4D). We also mapped several putative RpoS binding sequences within *flp*, *cupE,* and *rcpC* promoters (Fig. 4E). These results suggest that RpoS can also control the transcription of PprB-regulated genes in a PprB independent way, possibly through directly acting as sigma factors during transcriptional initiation.

### PprB overexpression enhances CCA in P. aeruginosa

We next investigated the possible effects of *pprB* upregulation on the physiology of *P. aeruginosa*. Overexpression of PprB was reported to result in a hyper-biofilm phenotype that was dependent on type IVb pili, the CupE fimbriae and the BapA adhesin (PA1874), but the exact mechanism by which this occurs remains unclear (22). During biofilm formation in flow chambers, bacteria are engaged in a dynamic process of growth and detachment, and the rate of bacterial growth and detachment within a biofilm are the two key factors that determine the resultant biomass. As the growth of *pprB* overexpression strain did not show any advantages to the wild type strain (Fig. S1), we speculated that the hyper biomass phenotype may be due to an enhanced cell-to-cell or cell-to-surface adhesion, both of which are thought to reduce the rate of detachment.

The CCA of bacteria was simply estimated by observing the formation of bacterial aggregates in shaking cultures at the logarithmic phase. Both the mean size and number of bacterial aggregates in the PprB overexpression strain are about twice that of wild type strain (Fig. 5A, Fig. 5B). Additionally, the established bacterial aggregates dispersed completely after a 30-min incubation with proteinase K at 37 °C (Fig. 5A). Thus, CCA was enhanced in the PprB overexpression strain, and the PprB regulated proteins may directly contribute to CCA. CCAs were then monitored in *flp*, *cupE*, and *bap* mutants under the background of PprB overexpression. The mean size of bacterial aggregates in these mutants displayed small differences from that in the wild type strain (Fig. 5C, grey bar). Whereas bacterial aggregates differ between strains in high-size regions, according to their size distribution curves (Fig. 5B). We counted the number of large aggregates whose size were bigger than 50 μm and determined that formation of large bacterial aggregates under a PprB overexpression background were, compared with wild type, (i) increased by 43% (P < 0.01) in the *flp* mutant, and (ii) reduced by 33% (P < 0.01) and 74% (P < 0.001) in the *cupE* and *bap* mutants, respectively (Fig. 5C). These results demonstrate that the Bap adhesin secretion system and CupE fimbriae are partially contribute to CCA, while the type IVb pili has a negative effect on CCA.

**Figure 5:**
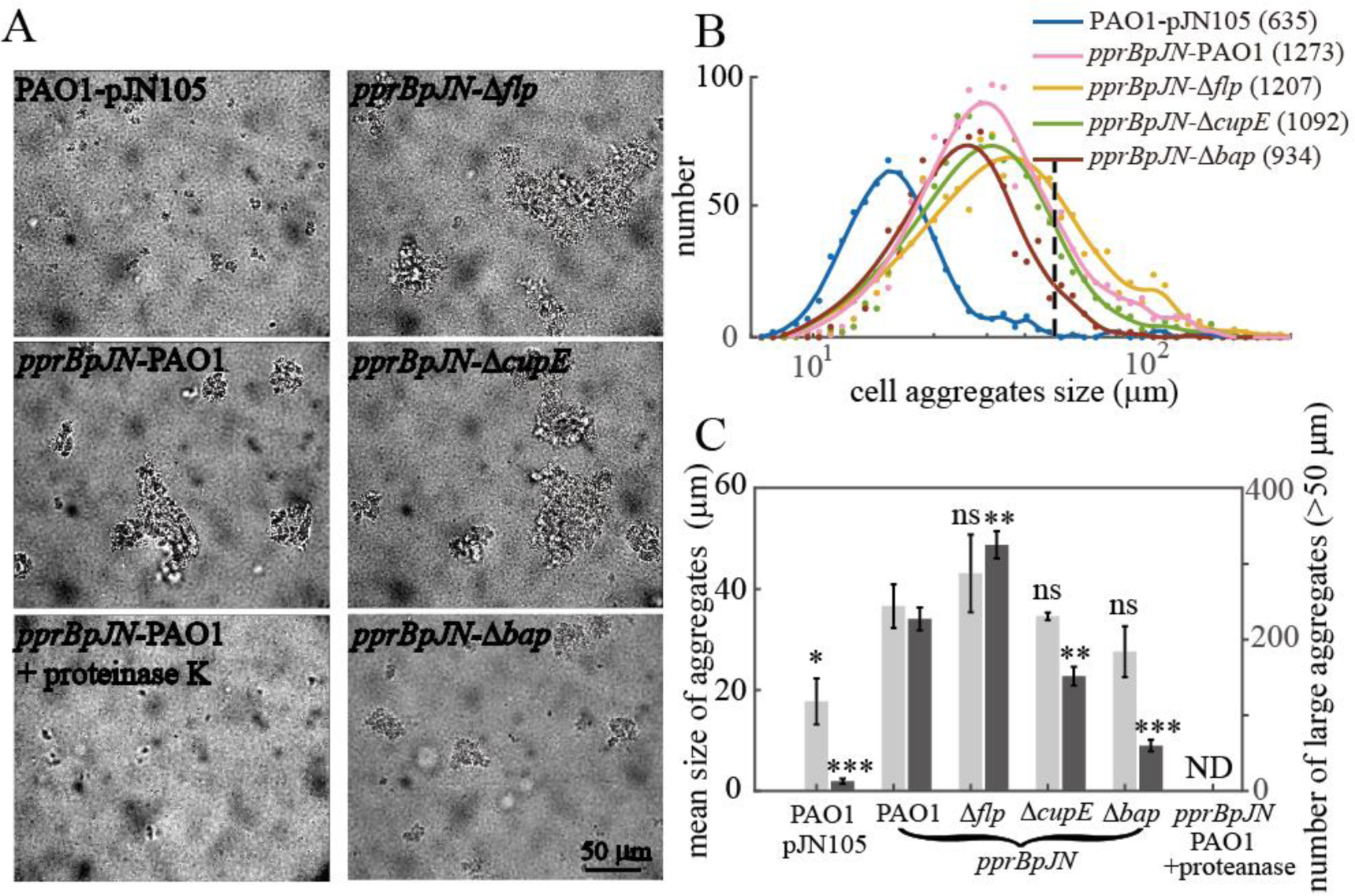
PprB overexpression enhances CCA in *P. aeruginosa*. (A), Bright field images of the logarithmic bacterial cultures of indicated strains. (B), Size distribution curves of cell aggregates of indicated strains, total numbers of cell aggregates counted for each strain are labelled in the brackets behind strain names. Data are presented on Logarithmic scale. The distribution curves are the smoothing result of original data points using Smoothing Spline method. The black dashed line indicates the position where cell aggregate size equals 50 μm. (C), Mean size of cell aggregates (light gray) and number of large aggregates (dark gray, cell aggregates size > 50 μm) in indicated strains. Mean aggregate sizes of samples are from three independent experiments and shown as the mean ± s.d. Error of big aggregates numbers are estimated from Poisson counts. ND, not detected. Statistical analysis used pairwise comparisons between corresponding data in the *pprBpJN*-PAO1 and other strains (t-test). *P < 0.05; **P < 0.01; ***P < 0.001; ns, non-significant.

Interestingly, cell clustering could only be observed when arabinose was added at the very start of bacteria inoculation. When arabinose was added at the logarithmic phase (OD_600_ ∼ 0.5), few clusters could be seen, and we also could not observe any clusters during the carbon deprivation experiment (in which cells can hardly grow). This phenomenon suggests that there is a currently unknown relationship between bacteria clustering and cell division.

### PprB overexpression enhances CSA in *P. aeruginosa*

The CSA of bacteria was measured using a microfluidic device. Bacterial cultures were injected into the device and stood for 20 minutes to enable initial adhesion. Then, we directly washed the microfluidic channel with shear stress of 70 Pa for 5 minutes, number of cells on the surface before and after shear stress were counted. The effect of *pprB* overexpression on CSA was investigated first. All strains were grown to the logarithmic phase before being injected to microfluidic channels. Fluidic shear eliminated most of the adhered cells in the wild type strain. By contrast, cells with PprB overexpression appeared to be largely unaffected, with only several cells with incomplete adhesion being eliminated (Fig. 6A), the remaining cells persisted sticking to the surface even when shear stress was increased to 1000 Pa. Under PprB overexpression background, fraction of remained cells after shear stress were, compared to wild type: (i) not changed in *flp* mutant; (ii) reduced by 27% (P < 0.05) and 79% (P < 0.001) in *cupE* and *bap* mutants respectively (Fig. 6B). Thus both the CupE fimbriae and the Bap secretion systems are involved in the enhanced CSA of the PprB overexpression strain.

**Figure 6:**
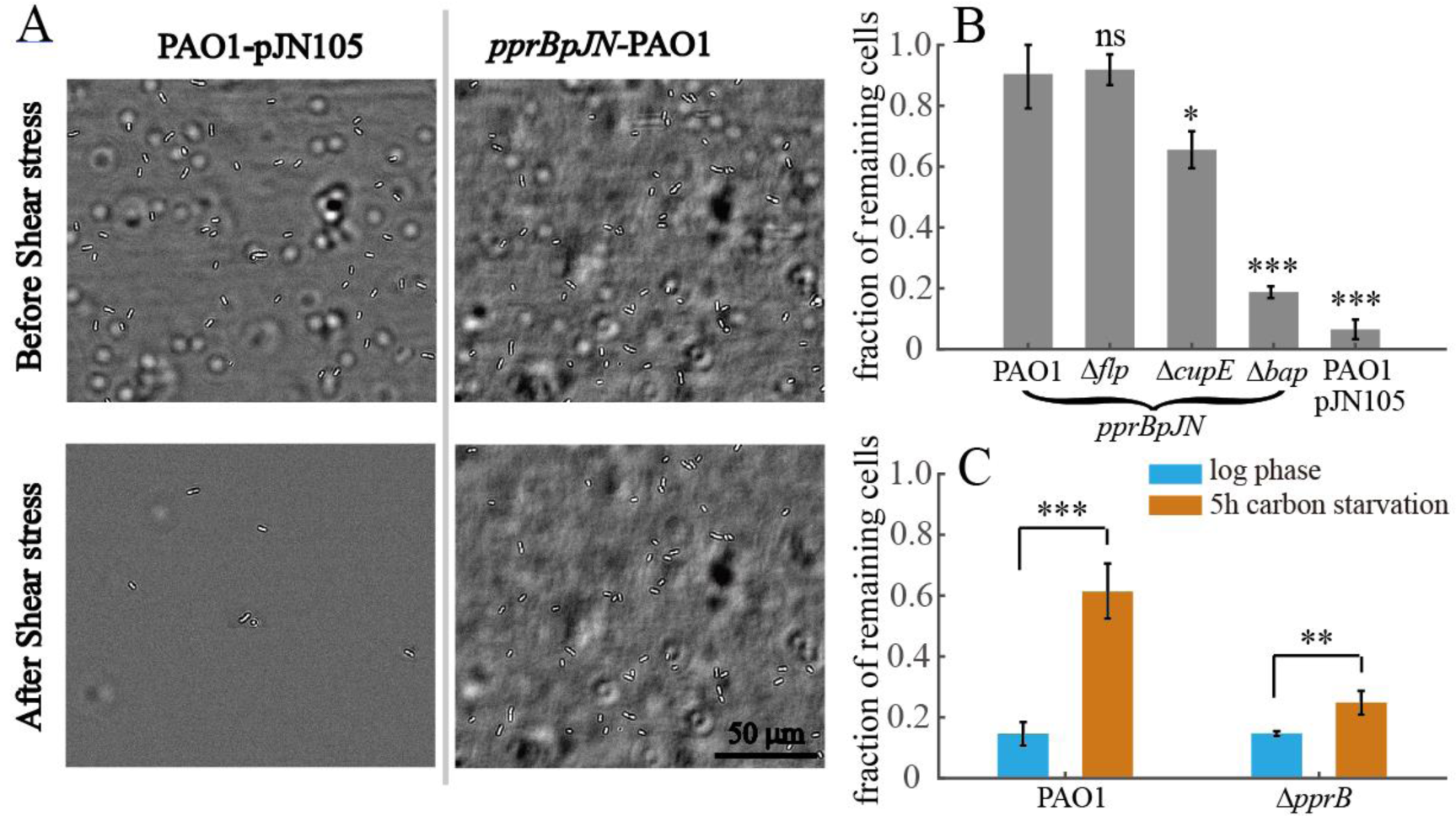
PprB overexpression enhances CSA in *P. aeruginosa*. (A), Bright field images of the wild type or PprB overexpression cells in microfluidic channels before and after exposing to 5-min shear stress (70 Pa). (B), Fraction of remaining cells on surface after exposing to 5-min shear stress (70 Pa). (C), Fraction of remaining cells on surface of wild type or *pprB* mutant strains after exposing to 5-min shear stress (70 Pa), cells were from the logarithmic phase or treated with 5-hour carbon starvation. Data in B and C are from three independent experiments and shown as the mean ± s.d. Statistical analysis used pairwise strain comparisons (t-test). *P < 0.05; **P < 0.01; ***P < 0.001; ns, non-significant.

We further monitored the CSA of the wild type and *pprB* mutant cells before and after carbon deprivation. As expected, the fraction of remaining cells in the wild type strain showed 4-fold (P < 0.001) increase upon CSS, in contrast to the 50% (P < 0.01) increase observed in the cells of the *pprB* mutant (Fig. 6C). Taken together, our results confirm that PprB overexpression can enhance bacterial CCA and CSA, which probably leads to the hyper biofilm phenotype.

Interestingly, although the type IVb pili is essential for the formation of the previously reported hyper-biofilm phenotype, this organelle showed no contribution to bacterial CSA (Fig. 6B) and showed a negative effect on bacterial CCA (Fig. 5C). The function of type IVb pili in biofilm formation remains unclear at this time.

### PprB negatively regulates the transcription of itself

Many transcriptional regulators in bacteria exhibit self-regulation activities, either positive or negative. PprB was reported to bind to the *pprB* promoter region(25), in which a putative PprB binding site (GGCTAATAC) was mapped based on a previously predicted PprB recognition consensus (Fig. 4A). The PprB binding site stands immediately downstream of the aforementioned RpoS site, suggesting a negative effect of PprB on *pprB* transcription due to the steric interaction between PprB and RNA polymerase. To verify this assumption, we measured the fluorescence of the *pprB* reporter during logarithmic phase or under carbon deprivation conditions in both the *pprB* mutant and overproducing strains. Under CSS, the activity of the *pprB* reporter in the *pprB* mutant strain is similar to that in the wild type strain, while in PApprB1 cells it was 20% (P < 0.001) of that in the wild type strain (Fig. 7A). Moreover, the expression of *PpprB-mut2* reporter whose PprB binding site was mutated (GGCTAATAC to GGCGGGTAC) was measured in the wild type and PApprB1 strains. In response to CSS, *PpprB-mut2* reporter in PApprB1 displayed 4-fold increase (P < 0.01), similar to the 5.5-fold increase (P < 0.01) found in the wild type strain (Fig. 7A). All these results confirm that *pprB* transcription is under a direct negative control of PprB. The model of CSS responses of PprB regulated genes through RpoS is presented (Fig. 7B).

**Figure 7:**
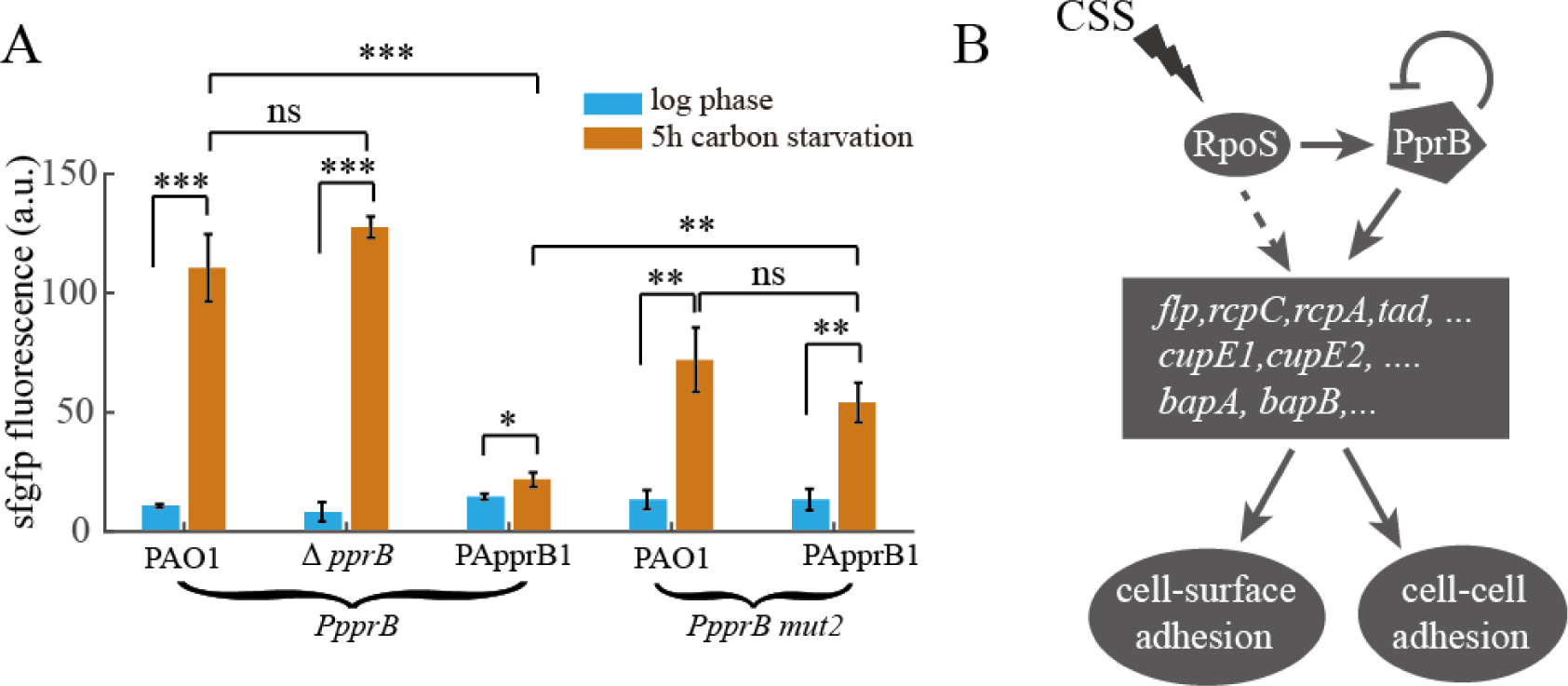
PprB negatively regulates the transcription of itself, and model of CSS responses of PprB-regulated genes through RpoS. (A), Expression values of *pprB* or *PpprB mut2* (PprB binding sequence GGCTAATAC was mutated to GGCGGGTAC) transcriptional reporters in wild type or *pprB* mutant or PApprB1 (PprB was constitutively overexpressed) strains at logarithmic phase or after 5-hour carbon deprivation. Data are from three independent experiments and shown as the mean ± s.d. (B), Schematic representation of the RpoS-PprB-Flp/CupE/Bap/Tad system and its signaling cascade to CSS. CSS induces the expression of PprB-regulated genes through triggering the expression of PprB. RpoS mediates the CSS signal induction of PprB transcription. Expression of PprB-regulated genes enhances bacterial CCA and CSA. PprB negatively regulates the transcription of itself. Statistical analysis used pairwise strain comparisons (t-test). *P < 0.05; **P < 0.01; ***P < 0.001.

## Discussion

The PprA-PprB two-component system has been studied for over 10 years, and PprB regulon containing multiple functional gene clusters was characterized several years ago. As it comes to the physiological role of PprB in bacteria, previous studies have mainly focused on the phenotypes of the *pprB* overexpression strain, which, compared with the wild type strain, have shown increased cell membrane permeability and aminoglycosides sensitivity, decreased cellular cytotoxicity and virulence in flies, and better biofilm formation (22, 23). However, very few results have been presented about the phenotypes of the *pprB* mutant strain, except one recent study that observed a compromised biofilm in this strain (24). The lack of knowledge about phenotypes of the *pprB* mutant is partially due to the fact that the signals and environmental conditions that may trigger the PprA-PprB system remain unclear. Generally, determining the external signals that can trigger a regulatory system is crucial to understand the regulatory logic and inward function of that system. In this paper, we provide evidence that the PprB-regulated genes could be induced via CSS. In particular, the induction of PprB regulated genes is dependent on the increased expression of PprB, rather than on the activation of the PprA kinase. We further demonstrate that the stress response sigma factor RpoS controls the induction of *pprB* transcription.

In many organisms, the small-molecule alarmone (p)ppGpp is the main effector of the stress response that takes place during starvation (36). The (p)ppGpp Synthases RelA can sense the inability of tRNA aminoacylation during carbon starvation and translate the carbon starvation signal to the synthesis of intracellular (p)ppGpp (37). RelA-dependent (p)ppGpp accumulation was also demonstrated in *S. suis* under CSS (38). In *E. coli,* (p)ppGpp positively affects the intracellular level and function of RpoS through the multifaceted regulation of transcription, translation, proteolysis, and activity (39), thereby tying the CSS signal to the response of the RpoS regulon. As most of the genes in the (p)ppGpp-RpoS system of *E. coli* can also be found in the *P. aeruginosa* genome, it is possible that the RpoS-dependent *pprB* transcriptional response observed in this study was achieved through the same (p)ppGpp-RelA stress-sensing mechanism. In addition, CSS is not the only signal that can induce PprB expression. Nitrogen starvation stress and acetate stress, two other signals that can trigger the RpoS stress-response system, also induce the transcription of *pprB* (Fig. 4C). Thus, signals facilitating the accumulation of intracellular RpoS are probably the signals that can activate the expression of PprB and PprB-regulated genes.

PprA was previously reported to be the cognate kinase for PprB (23). However, PprB is still active in the *pprA* mutant strain, according to the fact that *pprA* knockout failed to eliminate or reduce the CSS response of PprB-regulated genes. One reason could be that the regulatory activity of PprB is independent of PprB phosphorylation. This is contrary to our knowledge of two-component systems (40–42), thus the possibility seems unlikely. An alternative explanation is that there are other kinases which are responsible for PprB phosphorylation, this situation allows PprB to respond to other kinds of signals in addition to the RpoS related stress signals. Unfortunately, we have not found any kinase for PprB till now, a kinase-screening investigation in *P. aeruginosa* is needed in the future.

According to evolutionary theory, the induction of genes under a specific condition should be beneficial for the bacteria, whether through improved fitness or from enhanced competitive advantage over other organisms. We monitored the fitness of the wild type and *pprB* mutant strains under CSS in shaking cultures using the colony-forming unit method. Contrary to our expectation, the *pprB* mutant exhibited better fitness than the wild type strain did (Fig. 8A). Since biofilm is considered to be the natural form of existence of *P. aeruginosa,* and according to the previously found biological filtration effect of biofilm (43), cells in the deep inner regions of biofilm may encounter CSS as the biofilm grows and thickens. We detected the expression of both *pprB* and PprB regulated genes in *P. aeruginosa* colony biofilms. All the observed genes displayed thorough induction after 72 hours of incubation (Fig. 8B), indicating the probable involvement of PprB in biofilm development. Considering the enhanced cell-cell and cell-surface adhesions found in the PprB-overexpressed strain, and that the *pprB* mutant strain showed a reduced biofilm formation (24), this CSS-RpoS-PprB-BapA/Flp/CupE/Tad signaling pathway may help reinforce the structure of *P. aeruginosa* biofilms.

**Figure 8:**
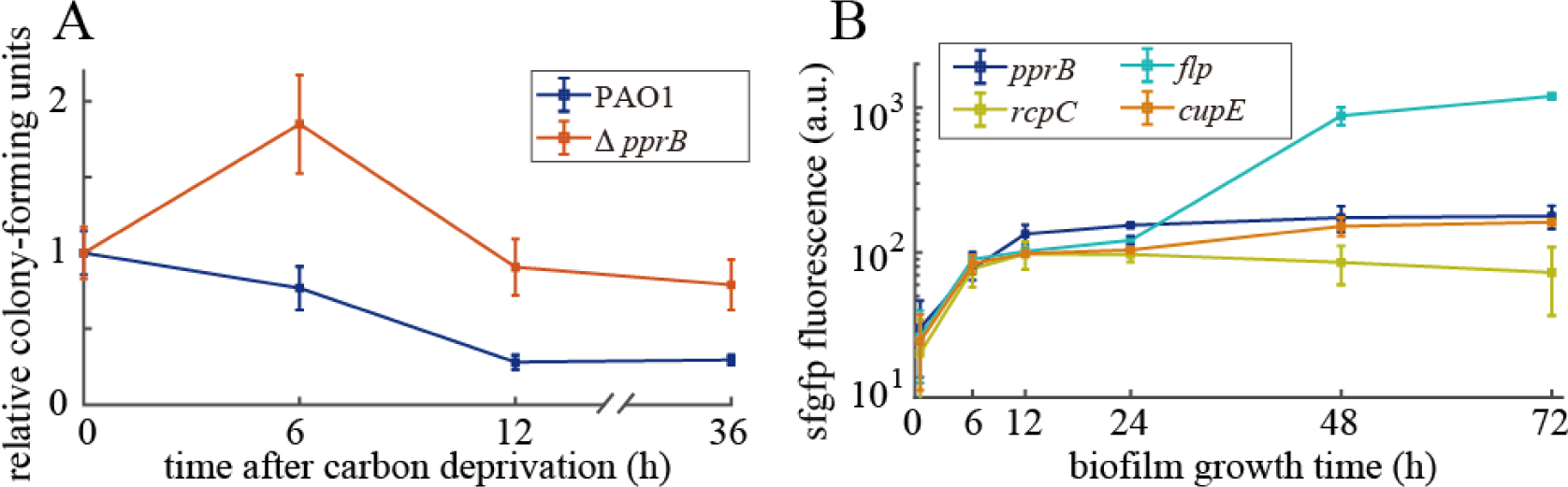
(A), Relative colony-forming units of wild type and *pprB* mutant cells after carbon deprivation for 0, 6, 12 and 36 hours in shaking conditions at 37°C. Colony-forming unit data of each strain were normalized by data at 0 hour. (B), Time dependent expression curves of *pprB, flp, rcpC or cupE* genes in the wild type cells grown in colony biofilms at 37°C. Data are from three independent experiments and shown as the mean ± s.d.

## Materials and Methods

### Bacterial strains and growth conditions

The strains and plasmids used in this study are listed in Table 1. Unless otherwise stated, cells were grown in FAB minimal media (44) supplemented with 30 mM sodium succinate (FABS) or other carbon sources (i.e., 30 mM sodium glutamate, 30 mM glucose, 10 mM alpha-Ketoglutaric acid (alfa-KG), 30 mM sodium citrate, 30 mM aspartic acid, or 30 mM sodium acetate) at 37 °C. To prevent plasmid loss, 30 μg/mL gentamycin was added to media for cultivation of the strains containing transcriptional reporter plasmids or pJN105-derivative vectors. LB media was used throughout the DNA cloning experiments. The *Escherichia coli* Top10 strain was used for standard genetic manipulations.

**Table 1:** Strains and Plasmids used in this study

| Strains or plasmids | description | origin |
| --- | --- | --- |
| ***E. coli* strain** |  |  |
| Top10 | F<sup>-</sup>, *mcrA*, (*mrr*; *hsdRMS*–*mcrBC*), 80*lacZ* M15 *lacX74*,<br>*recA1*, *araD139*, (*ara*–*leu*)7697, *galU*, *galK*, *rpsL*(Str<sup>R</sup>),<br>*endA1*, *nupG* | Invitrogen |
| ***P. aeruginosa* strains** |  |  |
| PAO1 | Wild-type strain | J.D. Shrout |
| $\Delta pprB$ | nonpolar *pprB* deletion in PAO1 | This study |
| $\Delta pprA$ | nonpolar *pprA* deletion in PAO1 | This study |
| $\Delta rpoS$ | nonpolar *rpoS* deletion in PAO1 | Kangming<br>Duan group |
| $\Delta flp$ | nonpolar *flp* deletion in PAO1 | This study |
| $\Delta cupE$ | nonpolar *cupE* deletion in PAO1 | This study |
| $\Delta bapA$ | nonpolar *bapA* deletion in PAO1 | This study |
| $\Delta lasR\Delta rhlR$ | nonpolar *lasR* and *rhlR* deletion in PAO1 | J.D. Shrout |
| $\Delta cbrA\Delta cbrB$ | nonpolar *cbrA* and *cbrB* deletion in PAO1 | This study |
| PApprB1 | $\Delta pprB$, *araC*-P<sub>BAD</sub>-*pprB*-miniTn7 | This study |
| PApprB2 | $\Delta rpoS$, *araC*-P<sub>BAD</sub>-*pprB*-miniTn7 | This study |
| PArpoS | $\Delta rpoS$, *PrpoS*-*rpoS*-miniTn7 | This study |
| PAO1 pJN105 | PAO1 strain containing pJN105 void vector, Gm<sup>R</sup> | This study |
| *pprB*pJN-PAO1 | PAO1 strain containing *pprB*-pJN105, *pprB* expression under | This study |
|  | the control of arabinose concentration , Gm<sup>R</sup> |  |
| *pprBpJN-Δflp* | *Δflp* strain containing *pprB*-pJN105, Gm<sup>R</sup> | This study |
| *pprBpJN-ΔcupE* | *ΔcupE* strain containing *pprB*-pJN105, Gm<sup>R</sup> | This study |
| *pprBpJN-ΔbapA* | *ΔbapA* strain containing *pprB*-pJN105, Gm<sup>R</sup> | This study |
| **Plasmids** |  |  |
| PUCPgfps | Cloning vector for transcriptional reporter,<br>*RNAseIII*-RBS2-*sfgfp*-T<sub>0</sub>T<sub>1</sub>-J23102-RBS2-*cyofp*-T-PUCP20,<br>Gm<sup>R</sup> | This study |
| *Pflp*-PUCPgfps | Transcriptional reporter plasmid of *flp*, Gm<sup>R</sup> | This study |
| *PcupE*-PUCPgfps | Transcriptional reporter plasmid of *cupE1* , Gm<sup>R</sup> | This study |
| *PrcpC*-PUCPgfps | Transcriptional reporter plasmid of *rcpC*, Gm<sup>R</sup> | This study |
| *PpprB*-PUCPgfps | Transcriptional reporter plasmid of *pprB*, Gm<sup>R</sup> | This study |
| *PpprB mut1* | Transcriptional reporter plasmid of *pprB*, with RpoS binding<br>site mutated from CTATATG to GGGTATG, Gm<sup>R</sup> | This study |
| *PpprB mut2* | Transcriptional reporter plasmid of *pprB*, with PprB binding<br>site mutated from GGCTAATAC to GGCGGGTAC, Gm<sup>R</sup> | This study |
| pex18ap | oriT<sup>+</sup>sacB<sup>+</sup>; gene replacement vector with MCS from<br>pUC18; Ap<sup>R</sup> | (46) |
| pex18gm | oriT<sup>+</sup>sacB<sup>+</sup>; gene replacement vector with MCS from<br>pUC18; Gm<sup>R</sup> | (46) |
| pFLP2 | sacB<sup>+</sup>; Flp recombinase-expressing bhr vector; Ap<sup>R</sup> | (46) |
| *flp*-gen-pex18ap | In-frame deletion of *flp* cloned into HindIII/XbaI sites of | This study |
|  | pex18ap; Ap<sup>R</sup>, Gm<sup>R</sup> |  |
| *pprB*-gen-pex18ap | In-frame deletion of *pprB* cloned into HindIII/XbaI sites of pex18ap; Ap<sup>R</sup>, Gm<sup>R</sup> | This study |
| *cupE*-pex18gm | In-frame deletion of *cupE* operon (*cupE1-cupE6*) cloned into pex18gm; Gm<sup>R</sup> | This study |
| *bapA*-pex18gm | In-frame deletion of *bap* operon (*bapA-bapD*) cloned into pex18gm; Gm<sup>R</sup> | This study |
| *pprA*-pex18gm | In-frame deletion of *flp* cloned into pex18gm; Gm<sup>R</sup> | This study |
| *araC*-P<sub>BAD</sub>-*pprB*-mi<br>niTn7 | *pprB* complementary plasmid for chromosomal insertion at attTn7 site, *pprB* expression is controlled by P<sub>BAD</sub> promoter | This study |
| *P<sub>rpoS</sub>*-*rpoS*-miniTn7 | *rpoS* complementary plasmid for chromosomal insertion at attTn7 site, *rpoS* expression is controlled by its own promoter | This study |
| *pprB*-pJN105 | *pprB* overexpression vector in pJN105, *pprB* expression is controlled by P<sub>BAD</sub> promoter | This study |

### Carbon deprivation experiment of transcriptional reporter strains

Overnight cultures of *P. aeruginosa* strains in FABS supplemented with 30 mg/mL gentamycin (FABSgen) were 100× diluted and grown to the logarithmic phase in FABSgen media. Cells were then harvested and washed once with FAB and resuspended in FAB + 30 μg/mL gentamycin (FABgen). Next, the suspensions were cultivated for a further 5 hours with shaking. For carbon deprivation in PApprB1 and PApprB2 strains, overnight cultures were diluted 100× in FABSgen + 0.4% (wt/vol) L-arabinose and grown to the logarithmic phase before following the same procedures noted above.

### Acetate and nitrogen starvation stress experiment

Overnight cultures of *pprB* transcriptional reporter strains in FABS supplemented with 30 μg/mL gentamycin (FABSgen) were 100× diluted and grown to the logarithmic phase in FABSgen media. For 200 mM acetate stress experiment, cells were then 100× diluted into FABSgen + 200 mM acetate and cultivated for 5 hours with shaking. Sfgfp fluorescence of cells was then measured by microscopy as mentioned below. For nitrogen starvation experiment, cells were washed once with and resuspended in the FABSgen medium without ammonium sulfate, and cultivated for 5 hours with shaking before use.

### Construction of gene deletion or complementary mutants in *P. aeruginosa*

PCR was used to generate 1000 bp DNA fragments upstream (Up) or downstream (Dn) from the *pprA*, *pprB*, *flp*, *cupE*, and *bap* genes. The primer pairs are listed in Table S1. The Up and Dn DNA fragments for *pprA*, *cupE* and *bap* were ligated together using overlap extension PCR, and then inserted into the pex18gm vector via Gibson assembly. The recombinant plasmids were introduced into *P. aeruginosa* through electroporation, and the deletion mutants were obtained by double selection on LB agar supplemented with gentamycin (30 μg/mL) and NaCl-free LB agar containing 15% sucrose at 37 °C (45). The Up and Dn DNA fragments for *cbrAB*, *pprB,* and *flp* were digested and cloned into pex18ap at HindIII/XbaI site together with *aacC1*. The recombinant plasmids were electroporated into *P. aeruginosa*, deletion mutants were obtained by selection on LB agar supplemented with gentamycin (30 μg/mL) containing 5% sucrose at 37 °C. Then pFLP2 system was used to delete the aacC1 cassette (46). The miniTn7 system (47) was used to construct the complementary *pprB* and *rpoS* mutants in *P. aeruginosa*. PCR fragments of *pprB* coding sequences and *araC-*P_BAD_ were inserted into the miniTn7 vector via Gibson assembly, generating P_BAD_-*pprB*-Tn7. Then, PCR fragments of the *rpoS* coding sequence, together with *rpoS* promoter sequence, were inserted into miniTn7 vector via Gibson assembly, generating *P_rpoS_*-*rpoS*-Tn7. The resultant plasmids were introduced to the *pprB* and *rpoS* mutant strains through electroporation, and the transconjugants were selected on 1.5% LB agar plates supplemented with 30 μg/mL gentamycin. The gentamycin resistance cassette in the complementary strains was then deleted according to a standard protocol (46).

### Construction of transcriptional reporters in *P. aeruginosa*

The *sfgfp* fusion plasmid used to measure the promoter activity of multi genes is a derivative of the vector PUCP20, here named pUCPgfp. *sfgfp*, *cyofp* and terminator fragments were amplified using PCR and inserted together into pUCP20 via Gibson assembly, generating PUCPgfp. The resultant genetic organization was *RNAseIII*-RBS2-*sfgfp*-T_0_T_1_-J23102-RBS2-*cyofp*-T-PUCP20. To construct the transcriptional fusion plasmids, promoter regions of *pprB*, *flp*, *rcpC*, and *cupE* were amplified by PCR from the PAO1 genomic DNA. The primer sets are noted in Table S1. Next, each fragment was cloned into PUCPgfp right before the RNAseIII site via Gibson assembly. The reporter plasmids were introduced into *P. aeruginosa* through chemical transformation. To construct site mutation transcriptional reporters of *pprB*, the wild type *pprB* reporter plasmid was used as a template. Two fragments were amplified: the primer pairs P*pprB*-mut1-F/PUCP20-R and P*pprB*–mut1-R/PUCP20-F were used for RpoS binding site mutation, and P*pprB*-mut2-F/PUCP20-R and P*pprB*– mut2-R/PUCP20-F were used for PprB binding site mutation. Then, the fragments were ligated together via Gibson assembly, generating P*pprB*-mutRpoS-PUCPgfp (P*pprB* mut1-*sfgfp*) and P*pprB*-mutPprB-PUCPgfp (P*pprB* mut2-*sfgfp*) plasmids. These two plasmids were introduced into *P. aeruginosa* strains through electroporation. All transconjugants were selected on 1.5% LB agar plates supplemented with 30 μg/mL gentamycin.

### Imaging of single cells of different promoter reporter strains and data analysis

The bacterial culture samples were pipetted out and loaded on a 2% (wt/vol) agarose FAB pad. Then, the pad was flipped onto a 0.15 mm cover glass so that the bacteria were sandwiched and lay flat between the agarose pad and the cover glass. Fluorescent images were acquired by confocal microscopy (IX-81, Olympus), equipped with a 100× oil objective and an EMCCD camera (Andor iXon897). Twenty-five image fields of each sample were snapped, from which more than 500 cells were imaged. In each image field, two images were acquired, one SfGFP image and one CyOFP image. SfGFP and CyOFP were both excited using a 488 nm laser and the fluorescence were collected through two emission filters, sized at 524 ± 25nm and 607 ± 25 nm. Data analysis was conducted using an image processing algorithm coded using MATLAB. Cell contours were obtained from the CyOFP images, then the SfGFP fluorescence of cells was measured by counting the mean intensities within corresponding cell contours in the SfGFP images.

### RNA-Seq experiment

Six parallel samples (50 mL each) were prepared, in which the overnight culture of PAO1 was diluted 50× in FABS media and grown until the mid-log phase (OD∼0.6) at 37 °C under shaking conditions. Three samples were stored at −80 °C, while the remaining three samples were washed 3 times with FAB and finally resuspended in FAB of the initial volume. These suspension cultures were cultivated for a further 6 hours with shaking at 37 °C and stored at −80 °C. The RNA extraction and sequencing procedures were performed in Guangzhou Huayin Medical Laboratory Center. Libraries were sequenced on an illumina HiSeq 2000 machine.

### Aggregation Assay

Overnight cultures of the *pprB* overexpression (in pJN105) or wild type strains were diluted 100× in FABSgen media supplemented with 0.02% (wt/vol) L-arabinose and grown to the mid-log phase (OD ∼ 0.6) at 37 °C. To measure the size of bacterial aggregates, 200 μL of each bacterial suspension were transferred into a 4-channel-dish (D35C4-20-1-N, Cellvis) and left to stand for 10 minutes at room temperature. The bacterial aggregates were monitored under a bright-field microscope equipped with a 60× oil objective. 100 images containing at least 200 bacterial aggregates were obtained every time. Three parallel experiments were conducted for each sample. The size of aggregates were recorded using ImageJ software. For Proteinase treatment, 20 μL proteinase K (R7012, Tiangen) was added to 1 mL bacterial culture at logarithmic phase and incubated for 30 minutes at 37 °C.

### Microfluidic experiment

For the microfluidic experiment of the *pprB* overexpression (in pJN105) strains, the culture conditions were the same as those for the aggregation assay. For the microfluidic experiment of the PAO1 and *pprB* mutant strains, the culture condition was the same as those for the carbon starvation experiment for transcriptional reporter strains, without the addition of gentamycin. The microchip platform was fabricated with polydimethylsiloxane (PDMS, Sylgard 184, Dow Corning) using standard soft lithography methods (48). Wafers were coated with SU-8 photoresist (MicroChem Inc., Newton, MA, USA) to form film deposition of up to 20 μm. The mould contained three parallel microchannels (length, 3 cm; width, 300 μm; and height, 20 μm) and was firmly stuck to a heat-tolerant plastic tray. Ten milliliters of the PDMS mixture, consisting of cross-linker and prepolymer PDMS (1:10, wt/wt), were added into the tray and baked at 80 °C f or 2 hours. The structure was then treated with a plasma cleaner (3 min) and bonded to a glass slide (Thermo Fisher Scientific Inc; length, 55 mm; width, 24 mm; thickness, 0.17 mm). In total, 0.5 mL of bacterial culture was injected into the channel for each experiment. The FAB medium was in a 10-mL gas-tight syringe and fluid flow was driven by a syringe pump (Harvard Apparatus, Holliston, Phd2000).

### Colony-forming units measurement

*P. aeruginosa* cultures cultivated under CSS for 0, 6, 12, and 36 hours were diluted up to 5000-fold with FAB medium, and plated in triplicate onto LB agar plates. Colonies were counted after a 24-hour incubation at 37 °C.

### Colony biofilm experiment

*pprB, rcpC, flp,* and *cupE* reporter strains of wild type *P. aeruginosa* were grown to the logarithmic phase in FAB medium containing 30 mM succinate and 30 μg/mL gentamycin, then 2 μL bacterial culture were gently dropped onto 1.5% agar plates containing the same medium. After the liquid on the culture plate evaporated, plates were incubated upside down at 37 °C for 6, 12, 24, 48, and 72 hours. Cells were scratched from the surface of biofilm colonies and resuspended in FAB medium before undergoing fluorescence measurement via microscopy.

## ACKNOWLEDGMENTS

We thank Kangming Duan for providing the *rpoS* mutant strain. We thank J.D. Shrout for providing the PAO1 wild type and *lasRrhlR* mutant strains. This work was supported by The National Natural Science Foundation of China (31700087, 21774117 and 31700745) and the Fundamental Research Funds for the Central Universities (WK3450000003) supported this work.

We were responsible for performing the study as follows: conceptualization, Lei Ni and Fan Jin; methodology, Wenhui Chen, Congcong Wang, Aiguo Xia, Rongrong Zhang, Lei Ni; investigation, Congcong Wang, Lei Ni, Fan Jin; writing— original draft—Lei Ni; writing—review and editing—Shuai Yang, Fan Jin;

